# The influence of biofilm formation on carbapenem resistance in clinical *Klebsiella pneumoniae* infections: phenotype vs genome-wide analysis

**DOI:** 10.1101/2020.07.03.186130

**Authors:** Naveen Kumar Devanga Ragupathi, Dhiviya Prabaa Muthuirulandi Sethuvel, Hariharan Triplicane Dwarakanathan, Dhivya Murugan, Yamini Umashankar, Peter N. Monk, Esther Karunakaran, Balaji Veeraraghavan

**Author notes:** Corresponding author: Dr. Balaji Veeraraghavan, Professor, Department of Clinical Microbiology, Christian Medical College, Vellore – 632004.

## Abstract

*Klebsiella pneumoniae* is one of the leading causes of nosocomial infections. Carbapenem-resistant (CR) *K. pneumoniae* are on the rise in India. The biofilm forming ability of *K. pneumoni*ae further complicates patient management. There is still a knowledge gap on the association of biofilm formation with patient outcome and carbapenem susceptibility, which is investigated in the present study.

*K. pneumoniae* isolates from patients admitted in critical care units with catheters and ventilators were included. *K. pneumoniae* (*n* = 72) were tested for antimicrobial susceptibility as recommended by CLSI 2019 and subjected to 96-well microtitre plate biofilm formation assay. Based on optical density at 570 nm isolates were graded as strong, moderate and weak biofilm formers. Subset of strong biofilm formers were subjected to whole genome sequencing and a core genome phylogenetic analysis in comparison with global isolates were performed. Biofilm formation was compared for an association with the carbapenem susceptibility and with patient outcome. Statistical significance, correlations and graphical representation were performed using SPSS v23.0.

Phenotypic analyses showed a positive correlation between biofilm formation and carbapenem resistance. Planktonic cells observed to be susceptible *in vitro* exhibited higher MICs in biofilm structure. The biofilm forming ability had a significant association with the morbidity/mortality. Infections by stronger biofilm forming pathogens significantly (P<0.05) resulted in fewer ‘average days alive’ for the patient (3.33) in comparison to those negative for biofilms (11.33). Phylogenetic analysis including global isolates revealed the clear association of sequence types with genes for biofilm mechanism and carbapenem resistance. Carbapenemase genes were found specific to each clade. The known hypervirulent clone-ST23 with *wcaG*, *magA*, *rmpA*, *rmpA2* and *wzc* with a lack of mutation for hyper-capsulation might be poor biofilm formers. Interestingly, ST15, ST16, ST307 and ST258 – reported global high-risk clones were *wcaJ* negative indicating the high potential of biofilm forming capacity. Genes *wabG* and *treC* for CPS, *bcsA* and *pgaC* for adhesins, *luxS* for quorum sensing were common in all clades in addition to genes for aerobactin (*iutA*), allantoin (*allS*), type I and III fimbriae (*fimA*, *fimH*, *mrkD*) and pili (*pilQ*, *ecpA*).

This study is the first of its kind to compare genetic features of antimicrobial resistance with a spectrum covering most of the genetic factors for *K. pneumoniae* biofilm. These results highlight the importance of biofilm screening to effectively manage nosocomial infections by *K. pneumoniae*. Further, data obtained on epidemiology and associations of biofilm and antimicrobial resistance genetic factors will serve to enhance our understanding on biofilm mechanisms in *K. pneumoniae*.

## Introduction

*Klebsiella pneumoniae* remains one of the leading causes of nosocomial infections and declared by WHO as a “priority pathogen”. Emergence of antibiotic-resistant *K. pneumoniae* worldwide have become a global concern. The strains that produce extended-spectrum β-lactamases (ESBLs) are increasing and carbapenems have become the choice to treat infections caused by ESBL-producing strains. In recent times, carbapenem-resistant *K. pneumoniae* (CRKP) have been on the rise, leading to multi-drug resistance (MDR) and limiting treatment options. Emergence of MDR *K. pneumoniae* is a cause for current concern in many countries worldwide with a mortality rate of ~42% for CRKP^[1]^.

*K. pneumoniae* are also known to cause biofilm-mediated infections in most of the hospitalised patients; this, in addition to carbapenem resistance, complicates the treatment. Biofilm-mediated infections have remained under researched for decades and their actual significance in influencing the outcome of antimicrobial therapy is yet to be fully understood. They are complex mono or polymicrobial structures, which result in persistent prolonged infections that are difficult to treat and clear *in vivo*. Most result from indwelling medical devices.

Some of the virulence factors of *K. pneumoniae* include capsule polysaccharide, lipopolysaccharide, type 1 and type 3 fimbriae, outer membrane proteins and determinants for iron acquisition and nitrogen source usage^[2,3]^. These virulence factors are known to be essential for the pathogen to evade host immune system and for successful biofilm formation^[2,3]^. The biofilm-forming phenomenon in *K. pneumoniae* was first described in 1988^[4]^. Investigations have proven that bacteria can survive for prolonged periods on inanimate surfaces like clinical devices, which promotes nosocomial infections^[5]^. Biofilm formation thus supports the ability of the bacteria to adhere to the indwelling medical devices like catheters and ventilators, which is expected to be coated with host cellular factors *in situ*.

Biofilm formation mechanisms in clinical *K. pneumoniae* is reported to be mediated by a series of genetic factors, including allantoin (*allS*), aerobactin (*iutA*), type I (*fimA* and *fimH*) and type III fimbriae (*mrkA* and *mrkD*), polysaccharides and adhesins (*pgaA*, *pgaB*, *pgaC*, *bcsA*), capsular polysaccharide (CPS) (*wzc, cpsD, treC, wcaG, wabG, rmpA/A2, magA, k2a, wzyk2*), quorum sensing (QS) (*luxS*) and colonic acid (*wcaJ*)^[6–11]^. Though these genetic factors were individually studied, a collective approach on their analysis in a set of clinical isolates is still lacking.

Antimicrobial resistance (AMR) is a global problem but very limited data are available on biofilm formation and its association with AMR among clinical isolates of *K. pneumoniae*. This study tries to address the gap in understanding the effect and association of *K. pneumoniae* biofilm formation on antimicrobial resistance in nosocomial infections. In addition, the study provides information on the epidemiology of *K. pneumoniae* on their genetic make-up for biofilm formation and which possibly enables the success of high-risk clones globally.

## Materials and Methods

### Study isolates

A total of 72 *K. pneumoniae* isolates previously obtained from clinical samples blood and endotracheal aspirates from patients in ICU/high-dependency units at the Christian Medical College, Vellore, India between 2018-2019 were selected for the study. These isolates were subjected to antimicrobial susceptibility and biofilm formation analyses, and whole-genome sequencing (WGS).

### Antimicrobial Susceptibility Testing

#### Disc diffusion

Antimicrobial susceptibility testing was performed by the Kirby-Bauer method with amikacin (30 μg), chloramphenicol (30 μg), tetracycline (30 μg), gentamycin (10 μg), ciprofloxacin (5 μg), cefotaxime (30 μg), cefoxitin (30 μg), ceftazidime (30 μg), cefpodoxime (10 μg), piperacilllin-tazobactam (100/10 μg), cefoperazone-sulbactam (75/30), netilmicin (30 μg), imipenem (10 μg), meropenem (10 μg) and tigecycline (15 μg) according to guidelines suggested by CLSI M100-S29, 2019. Quality control strains used were *E. coli* ATCC 25922 for all antibiotics concurrently in all the batches. Tigecycline results were interpreted according to FDA criteria.

#### Minimum Inhibitory Concentration Testing (MIC)

MIC tests were performed for meropenem by the broth microdilution method. *E. coli* ATCC 25922 was used as quality control strain for MIC determination with the expected range of 0.008 – 0.06 μg/ml for meropenem. The interpretive criterion provided by CLSI 2019 for susceptible, intermediate and resistant strains were ≤4 μg/ml, 8 μg/ml and ≥16 μg/ml for meropenem.

### Screening for biofilm formation

#### Biofilm screening assay

The protocol used was slightly modified from the method described by Di Domenico et al^[12]^. About 5-10 colonies from fresh culture were inoculated in a 10 ml LB broth and incubated for 12-18 h at 37°C. Optical density (OD) was measured in a spectrophotometer (Shimadzu, Kyoto, Japan) at 625 nm and 0.05 OD_625_ cells were prepared by dilution in Mueller-Hinton broth (MHB) broth (1% glucose). 150 μl of prepared cells were inoculated into each well on a 96 well plate and incubated at 37°C for 24 h. After incubation, the medium was removed and the biofilm was washed with 200 μl distilled water twice. For staining, 200 μl of 0.1% (w/v) crystal violet stain was added and incubated for 10 min at RT. Stain was removed and wells were washed twice with distilled water. To dissolve the stain, 200 μl of 33% (v/v) glacial acetic acid was added and incubated for 5 min at RT. OD was read at 570 nm. The assay was performed in triplicates and a blank well containing only growth medium without cells were used as negative control for the calculation of biofilm formation efficiency. Mean and SD was calculated for statistical significance.

The OD_570_ values were used to compare and classify biofilm production semi-quantitatively. Accordingly, the cut-off OD (ODc) was calculated mean OD of negative control with three standard deviations. Biofilm production was classified as: OD < ODc = poor biofilm producer; ODc < OD ≤ 2 × ODc = weak biofilm producer; 2 × ODc < OD < 4 × ODc = moderate biofilm producer; and OD ≥ 4 × ODc = high biofilm producer.

#### Minimum biofilm eradication concentration (MBEC) estimation assay

An MBEC assay was performed for the study isolates using the method previously described by Chen et al.^[13]^ with slight modifications. Briefly, all isolates were grown in cation-adjusted MHB with 1% glucose at 37 °C overnight. Cultures were then adjusted for the OD to meet 0.5 McFarland standards (0.05 OD) using MHB. 150 μL adjusted inoculum was added to all the wells in a 96-well MBEC Assay^®^ bottom plate (Innovotech, AB, Canada), and the MBEC inoculator plate was inserted and incubated at 37 °C for 24 h to allow biofilm formation. After the production of biofilm in the inoculator plate, four pegs from it were taken using a sterile tool and tested for the density of biofilm formed. The pegs were put in 200 μl TSB medium with 1% Tween20 (rich medium) in wells A1 – A4, and 20 μl of inoculum from the suspension (planktonic cells) in the MBEC bottom plate (any four random wells) were inoculated in wells A5-A8 with 180 μl of rich medium. The plate was sonicated for 10 min at high power and then the suspension was serially diluted up to 10^−8^ dilutions. 20 μl of the diluted inoculum was plated on LB agar and incubated at 37 °C overnight. The rest of the inoculator plate with biofilm was inoculated into a 96-well plate containing various concentrations of the antibiotic in 200 μl MHB in duplicates. The plate setup was incubated at 37 °C for 24 h. Following incubation, the peg plate was removed and washed with sterile distilled water for 1 min and introduced in a 96-well bottom plate with 200 μl of TSB medium with (1% tween 20). The plate was sonicated for 10 min at high power. Each well was diluted up to 10^−4^ dilutions and 20 μl was plated in LB agar followed by incubation at 37 °C for overnight. The LB plates were recorded for CFU/ml and the MBEC values obtained.

### Whole genome sequencing (WGS)

Isolates were sequenced for analysis of carbapenem resistance and other genetic factors involved. Genomic DNA was extracted with QIAamp DNA mini kit (Qiagen, Hilden, Germany). Whole genome sequencing (WGS) was performed using Ion Torrent (PGM) sequencer with 400-bp read chemistry (Life Technologies) according to the manufacturer’s instructions. The data was assembled *de novo* using AssemblerSPAdes v5.0.0.0 embedded in Torrent suite server version 5.0.3. The sequence annotation was performed in NCBI Prokaryotic Genomes Automatic Annotation Pipeline (PGAAP) (https://www.ncbi.nlm.nih.gov/genome/annotation_prok/). Downstream analysis was done using the Center for Genomic Epidemiology (CGE) server (http://www.cbs.dtu.dk/services).

### Genome-wide association and phylogenetic analysis

In addition to the isolates sequenced in this study, complete genome sequences of 473 isolates with reported biofilm genotypes were selected and downloaded from NCBI for a global comparison of the study isolates. Accordingly, *K. pneumoniae* and *K. quasipneumoniae* were analysed separately for their phylogenetic relations. Fasta sequences were used to call core SNPs using Snippy v4.4.0 (https://github.com/tseemann/snippy) and recombinations were removed using Gubbins v2.0.0^[14]^. RAxML program was used to build the phylogenetic trees using the clean core SNP alignments generated from Gubbins.

Obtained 473 fasta sequences were annotated using Prokka v1.14.6 for further analysis^[15]^. The isolates were analysed using Roary v3.11.2^[16]^ with Mafft v7.467 to identify the core genes involved among the selected population of *K. pneumoniae*, which was used to study the genes involved in biofilm production. Sequences were then analysed using ABRicate v0.8.7 for their AMR genes and plasmids; MLST was identified using mlst v 2.18.0 algorithm (https://github.com/tseemann/mlst). Individual biofilm gene sequences were retrieved using a blast algorithm, which were further subjected to Snippy for generating a core alignment and the tree was built using RaxML. All trees generated in the study were visualised using iTOL v 5.5.1. Scoary v1.6.16 was used for genotypic association analysis with parameters ‘-c I EPW -p 0.1 0.05 -collapse’ by providing core genome alignment tree generated by Roary^[17]^.

This Whole Genome Shotgun project has been deposited at GenBank under the accession numbers JAAMFF00000000, JAALJL00000000, JAALJK00000000, JAAMFE00000000, JAAMFD00000000 and JAALJJ00000000.

### Statistical analysis

The statistical parameters in the study were calculated using SPSS 16.0, GraphPad Prism v8.2.0 and Microsoft Excel 2016 (Roselle, IL, USA).

## Results

### Antimicrobial resistance

Among the 72 isolates tested from patients under critical care, ~20% had carbapenem susceptibility, followed by minocycline and tigecycline, with 30% and 45% susceptibilities, respectively. In contrast, most of other tested antimicrobials had <20% susceptibility.

### Association of carbapenem resistance and clinical outcome with biofilm formation efficiency

The biofilm formation assay revealed that out of 72 isolates tested, 27.78% (*n* = 20) were strong biofilm producers and 30.56% (*n* = 22) were negative for biofilm production. In addition, strong biofilm forming pathogens exhibited comparatively higher carbapenem resistance compared to moderate, weak and negative biofilm formers suggesting a positive correlation between biofilm formation and carbapenem resistance (Table 1).

**Table 1:**
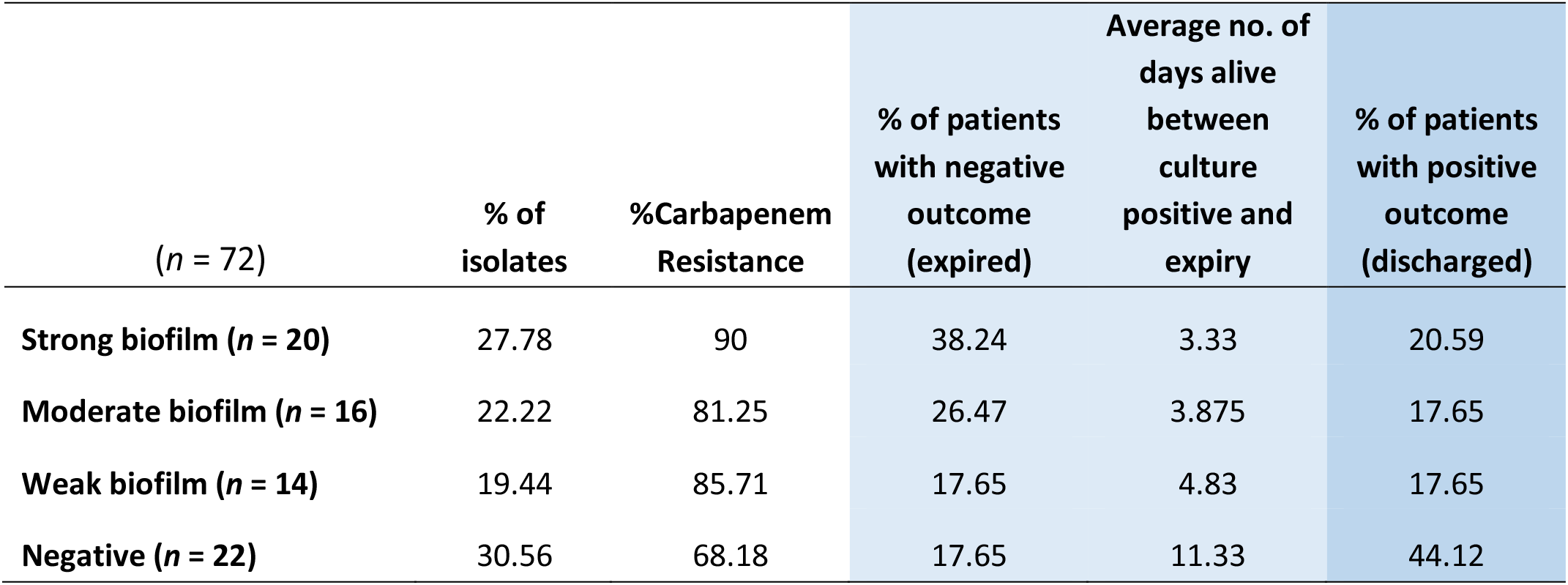
Showing distribution of morbidity, mortality and average days alive for patients with strong, moderate, weak and negative biofilm forming *K. pneumoniae* infections

Comparison of mortality and length of hospital stay from the onset of infection revealed the average ‘days alive’ of a patient for weak biofilm – 4.83, moderate biofilm – 3.875, strong biofilm forming pathogens – 3.33, when compared to 11.33 days for patients with isolates negative for biofilm production. This was statistically significant (p<0.05), showing that infection by stronger biofilm forming pathogens resulted in comparatively fewer days alive for the patient (Table 1). In addition, the number of patients expired were significantly higher (38.2%) in infections with strong biofilm forming *K. pneumoniae*, in comparison with biofilm negative *K. pneumoniae* (17.64%).

Interestingly, around 62% (*n* = 45) of the patients with *K. pneumoniae* were found co-infected with bacterial and yeast pathogens, with both monomicrobial and polymicrobial co-infections (Figure 1). Pathogens obtained in addition to *K. pneumoniae* from same patient were considered as co-infection. These included *P. aeruginosa* (*n* = 11), *Acinetobacter spp.* (*n* = 11), non-fermenting Gram-negative bacteria (NFGNB) (*n* = 4), *E. coli* (*n* = 5), *Klebsiella spp.* (*n* = 2), *Enterococcus spp.* (*n* = 5), *S. aureus* (*n* = 4), CoNS (*n* = 9), Group B *Streptococcus* (*n* = 2), *Aeromonas spp* (*n* = 1) and yeast (*n* = 12). There was no significant difference between the numbers in groups of strong, moderate, weak and negative biofilm producers.

**Figure 1.**
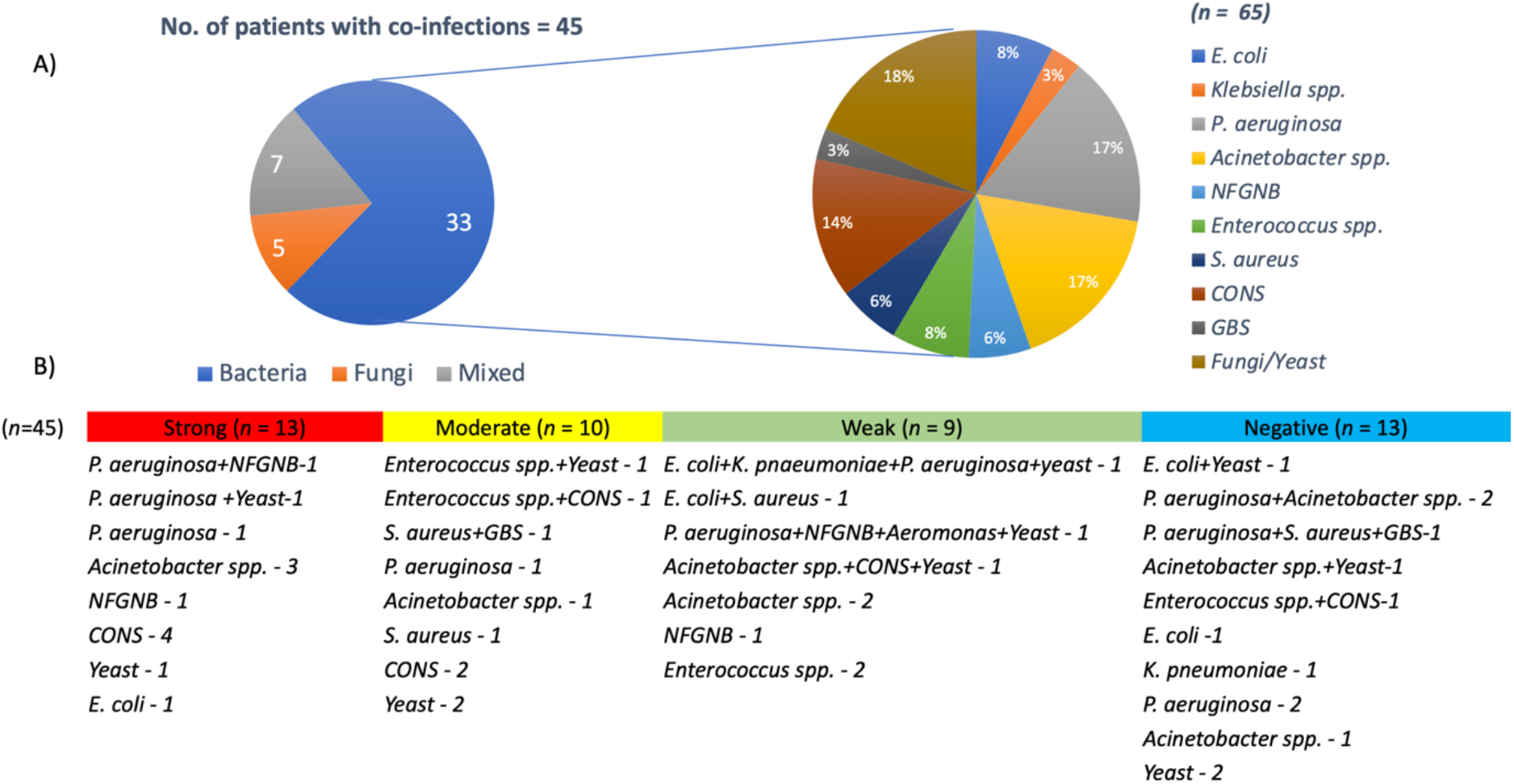
Pathogens co-infecting patients with *K. pneumoniae*. A) Of 45 *K. pneumoniae* infections, 33 were co-infected with bacteria, 5 with yeast and 7 were mixed infections. Also depicted is the number of each clinical pathogen among the bacterial co-infected population B) Numbers on each category with classification under strong, moderate, weak and negative biofilm forming *K. pneumoniae*.

### MBEC analysis

The MBEC of meropenem differed in comparison to MICs for the eight strong biofilm forming *K. pneumoniae* isolates tested (Table 2). The MBEC values for biofilms were higher than the MIC values estimated for planktonic cells. The quantitative log growth values (CFU/ml) of *K. pneumoniae* at various concentrations of meropenem were depicted in Figure 2.

**Table 2:**
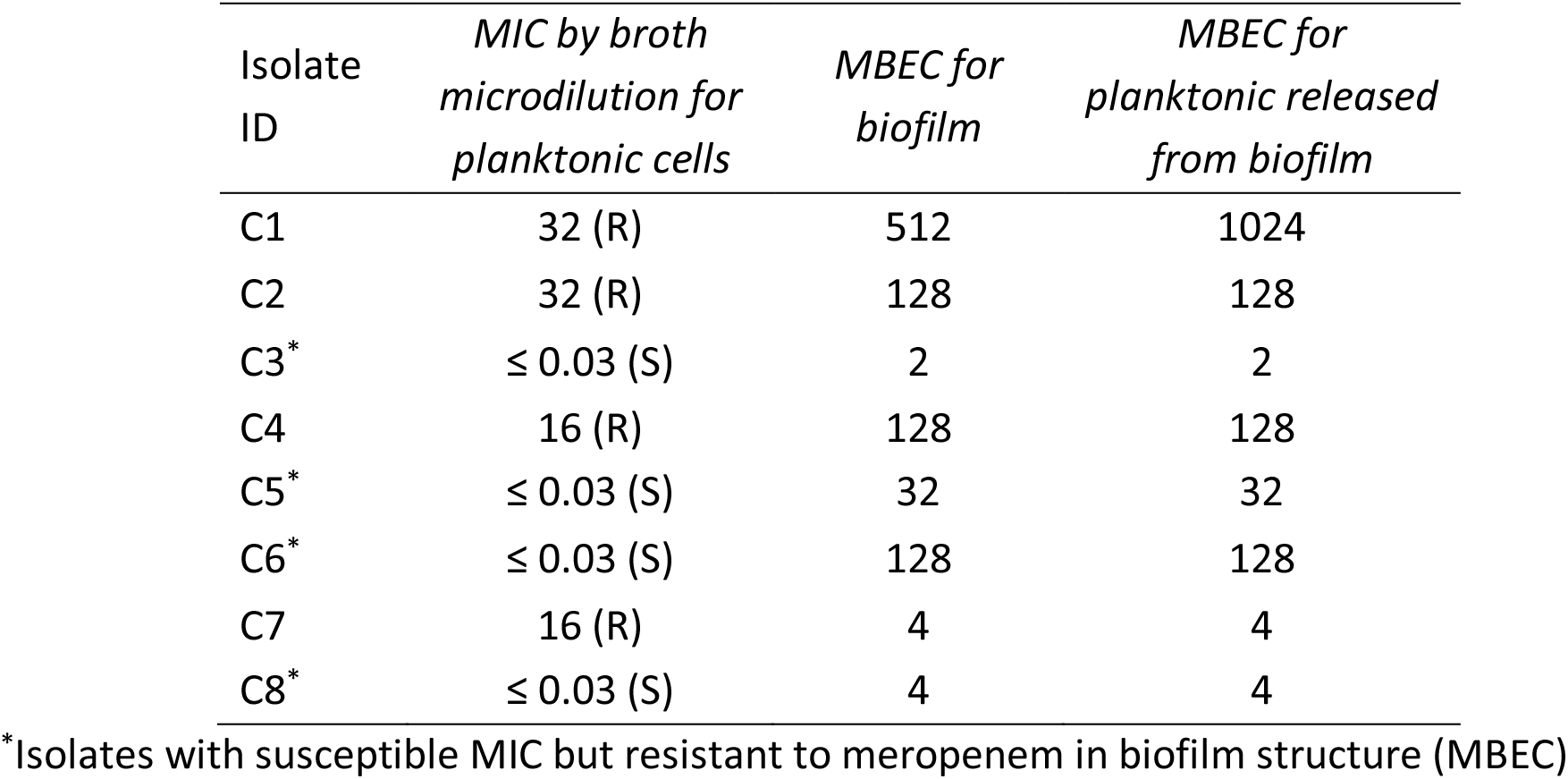
MBEC values of meropenem towards biofilm forming clinical *K. pneumoniae*

**Figure 2:**
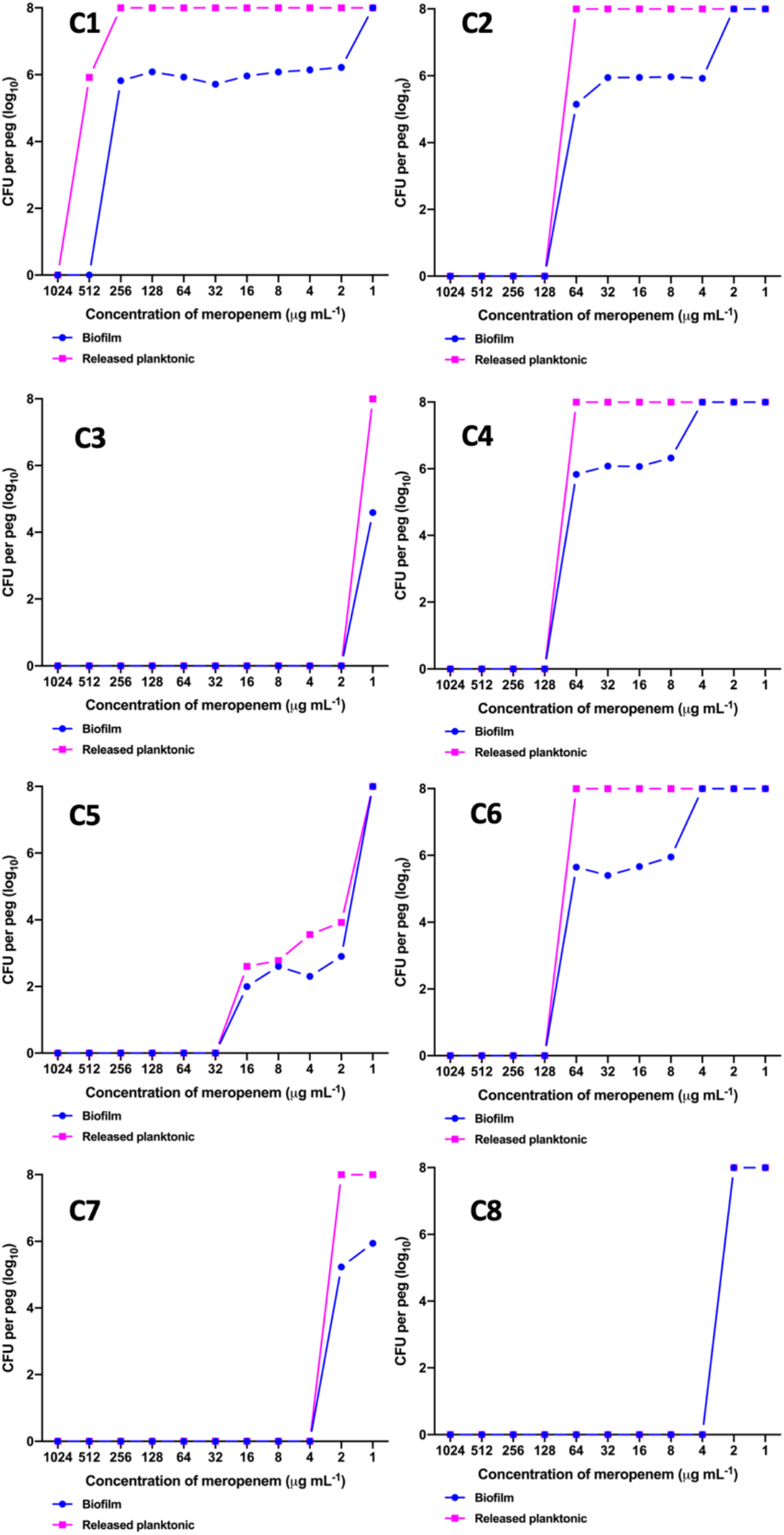
MBEC assay comparing the biofilm eradication efficiency of meropenem *in vitro* on *K. pneumoniae* with biofilm structure. X-axis indicates concentration of meropenem used for treatment, Y-axis indicates log_10_ CFU/ml of cells at respective concentrations. Data was visualised using GraphPad Prism v8.2.0. Biofilm – *K. pneumoniae* biofilm structure formed on pegs; Released planktonic – Cells released from the biofilm structure on pegs during meropenem treatment.

### Genome analysis of K. pneumoniae and K. quasipneumoniae

A representative sample of six strong biofilm forming *K. pneumoniae* were sequenced to ~60X coverage. WGS revealed three of these isolates to be *K. quasipneumoniae*. These were compared with global isolates obtained from NCBI for further analysis. Core genome analysis of 454 *K. pneumoniae* using Roary revealed 1129 core genes (in 99%), 1511 soft core (95-99%) and 3439 shell genes (15-95%), 36072 cloud genes (<15%), total 42151, whereas for 25 *K. quasipneumoniae*, 4931 core genes, 1394 shell genes, and 6325 total genes were found. Total SNPS included for phylogenetic analysis were 180537 for *K. pneumoniae* and 197628 in *K. quasipneumoniae*.

### Association of global clones with genes responsible for biofilm mechanism and carbapenem resistance

Among both *K. pneumoniae* and *K. quasipneumoniae*, genes responsible for biofilm formation, *allS* (aerobactin), *iutA* (allantoin), type 1 fimbriae (*fimA, fimH*), type III fimbriae (*mrkD*), pili (*pilQ, ecpA*), adhesins/polysaccharides (*pgaA, pgaB, pgaC, bcsA*), CPS (*wzc, cpsD, treC, wcaG, wabG, rmpA/A2, magA, k2a, wzyk2*), QS (*luxS*), colonic acid-mucoid (*wcaJ*) were screened and presented in figures 3 and 4.

**Figure 3:**
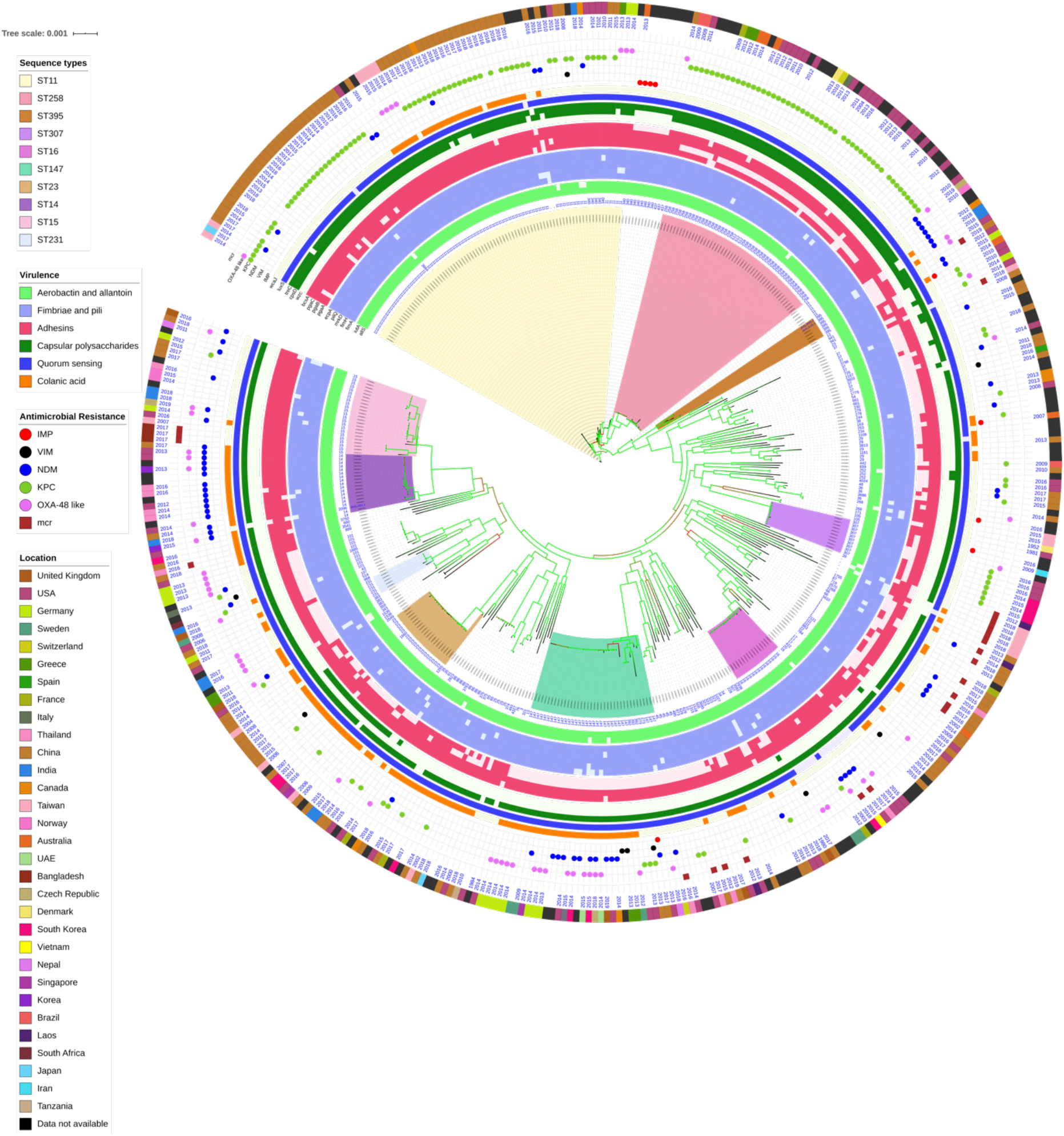
Core genome phylogeny of strong biofilm forming clinical *K. pneumoniae* in comparison to global genomes for identifying similarities in biofilm virulome and resistome. The color of branch leaves indicate the bootstrap values, green indicating high; red indicating low. Sequence types were marked in color ranges; innermost circle represents biofilm virulome (*allS, iutA, fimA, fimH, mrkD, pilQ, ecpA, pgaABC, bcsA, wzc, cpsD, treC, wcaG, wabG, rmpA/A2, magA, k2a, wzyk2, luzS, wcaJ*) followed by resistome (*bla*_IMP_, *bla*_VIM_, *bla*_NDM_, *bla*_KPC_, *bla*_OXA-48_ like, *mcr*) as shape plots. Location of each isolate has been given in the outermost ring as colour strip.

**Figure 4:**
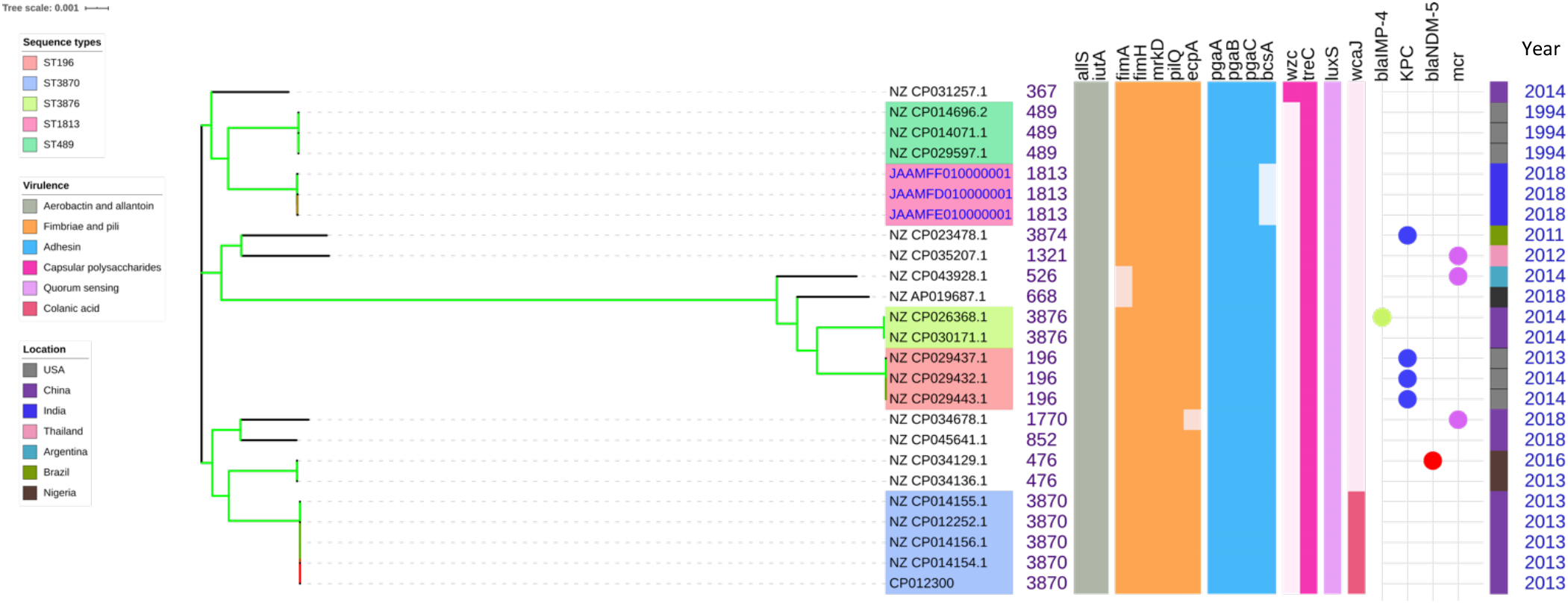
Comparison of *K. quasipneumoniae* clinical isolates to the global complete genomes indicating the biofilm virulome and resistome. The color of branch leaves indicate the bootstrap values, green indicating high; red indicating low. Sequence types were marked in color ranges followed by biofilm virulome (*allS, iutA, fimA, fimH, mrkD, pilQ, ecpA, pgaABC, bcsA, wzc, cpsD, treC, wcaG, wabG, rmpA/A2, magA, k2a, wzyk2, luzS, wcaJ*) and resistome (*bla*_IMP_, *bla*_VIM_, *bla*_NDM_, *bla*_KPC_, *bla*_OXA-48_ like, *mcr*). Colour strips indicate the country of the isolate.

In *K. pneumoniae*, the comparison of sequence type (STs) correlated well with the biofilm forming virulence genes, AMR genes and country of isolation. ST11 and ST258 were the major clades harbouring *bla*_KPC_ for carbapenem resistance. These clades were predominantly observed in the Americas, Europe and China. ST11 were predominantly seen in China, with only small numbers in USA, Europe and Taiwan. Isolates from Europe and Taiwan alone harboured *bla*_OXA-48_ like carbapenemases instead of *bla*_KPC_. A fraction of isolates from China harboured *bla*_NDM_ either alone or in addition to *bla*_KPC_. In contrast, ST258 was a strict *bla*_KPC_ clone and did not harbour any other carbapenemases. ST258 was predominantly found in USA followed by fewer isolates in Europe and Australia.

ST11 harboured most of the genes required for the process of biofilm formation, *allS, iutA, fimA, fimH, mrkD, pilQ, ecpA, pgaA, pgaB, pgaC, bcsA, cpsD*, *treC* and *wabG*. ST258 also carried the same set of genes for biofilm except *pgaB*, which was clearly missing in ST258. Both ST11 and ST258 did not harbour *wzc, wcaG, magA, k2a* and *wzyk2* for CPS. *rmpA* and *rmpA2* were found in few of the ST11 isolates (*n* = 20) from China. Almost 50% of ST11 from China lacked *wcaJ* known for colanic acid production, while none of ST258 isolate carried *wcaJ*.

ST395 was typically missing *pgaA* and *pgaB* genes for polysaccharide production. They were distributed in the USA (*n* = 2), India (*n* = 2), China (*n* = 1) and Germany (*n* = 1). One of the USA isolates carried *k2a* and *wzyk2* genes for K2 serotype. Two of the study isolates from India lacked *bcsA* and *treC* in addition to *pgaA* and *pgaB*. These isolates harboured only one gene for adhesin (*pgaC*), while carrying *cpsD*, *treC* and *wabG* for CPS. Except two isolates from India (C1) and USA, none of ST395 carried *wcaJ*. Four out of six ST395 isolates harboured *bla*_NDM_, with one Indian isolate harbouring both *bla*_NDM_ and *bla*_OXA-232_. One of the two isolates lacking *bla*_NDM_ was found harbouring *bla*_OXA-48_ like. Interestingly, one isolate from China that completely lacked carbapenemases carried an *mcr* gene.

ST307, which is again a *bla*_KPC_ predominant clone, was observed in the USA and South Korea. This clone lacked the *pgaB* gene for adhesin but carried *treC* and *wabG* for CPS. ST16 is an another reported MDR clone, observed in USA and Europe, with one isolate each in Thailand, Vietnam and South Korea. ST16 had a mixture of *bla*_NDM_ and *bla*_OXA-48_ like carbapenemases. Interestingly, one isolate each from Thailand and Vietnam harboured *mcr* in addition to *bla*_NDM_. ST16 harboured only *treC* and *wabG* for CPS but harboured all other screened genes for type I and III fimbriae, adhesins, aerobactin and allantoin.

ST147 is a reported *K. pneumoniae* global clone and was observed across all countries. However, based on the geographical occurrence, the carbapenemase genes carried differed. ST147 *K. pneumoniae* from Germany (*n* = 8), UAE (*n* = 2), Pakistan (n = 1) and Nepal (*n* = 1) harboured *bla*_OXA-48_ like, while one isolate from Greece harboured *bla*_KPC_. *bla*_NDM_ was observed in ST147 from Singapore, Switzerland, UK and Canada. In addition, a combination of *bla*_NDM_ and *bla*_OXA-48_ was observed in South Korea (*n* = 2) and Czech Republic (*n* = 1).

ST147 from USA harboured both *bla*_NDM_ and *bla*_KPC_, while two isolates from China harboured each with *bla*_KPC_+*bla*_VIM_ and *bla*_NDM_+*bla*_IMP_. One ST147 isolate from Thailand carried an *mcr* gene for colistin resistance. ST147 from Sweden (*n* = 2) did not harbour any carbapenemases in 2009, later in 2012 an isolate found acquired *bla*_OXA-48_ like. ST147 lacked *pgaA* and *pgaB* genes for adhesin, instead harboured *pgaC*, *bcsA*. For CPS ST147 carried only *treC* and *wabG*, while carrying *wcaJ* for colanic acid.

ST23 was mostly found in China, with a few isolates identified in USA, South Korea and India. Few isolates from China and USA alone carried *bla*_KPC_, whereas one isolate from China and India carried *bla*_VIM_ and *bla*_OXA-48_ like, respectively. ST23 harboured all screened genes for aerobactin, allantoin, fimbriae, pili, and adhesins except *pgaA*. For CPS, they exclusively carried *wzc* and *magA* gene in addition to *wabG* and *treC*, which was not seen in any related/unrelated STs. Also, *wcaG, rmpA*/*A2* were seen majorly in ST23 with very few reports in other STs. Since these were the only ST harbouring *wzc* genes for CPS, they were further analysed for their efficiency in producing hyper-capsulation by carrying a 565 glycine-to-serine substitution in *wzc* genes. The analysis revealed that none of the isolates were mutated at 565 glycine-to-serine position, known for hyper-capsulation.

ST231 and ST101 are *bla*_OXA-48_ carrying clones thought to be split from a single parent clade but still appear to converge in the core gene characteristics that are evident in their biofilm and AMR gene profiles. In this study population, ST231 was identified in India and the USA. Interestingly one isolate from ST101 carried *bla*_KPC_ from India. Both ST231 and ST101 lacked *pgaA* and *pgaB* genes for adhesins and carried only *treC* and *wabG* for CPS. Most of ST231 and ST101 lacked *wcaJ* for colanic acid. Similarly, ST14 and ST15 were diverged from a single parent clade found across most of the countries, where ST14 carried *bla*_NDM_ in almost all isolates (*n* = 17) in addition to *bla*_OXA-48_ like in few (*n* = 5). Whereas, ST15 started to lose *bla*_NDM_ that only carried *bla*_NDM_ gene in 6 out of 23 isolates. ST14 differed from ST15 in their CPS gene profile and *wcaJ*. ST14 harboured *k2a* and *wzyk* responsible for K2 serotype along with *wcaJ* gene. In addition, the sequenced study isolate C4 carried *rmpA* and *rmpA2*. Among ST15, only one isolate from USA carried *k2a*, *wzyk2* and *wcaJ* while others lacked these genes. Five out of 23 isolates harboured *rmpA* and *rmpA2*.

*luxS* responsible for QS in *K. pneumoniae* was present in almost all isolates irrespective of STs and countries. *mrkD*, and *treC* were further analysed for differences in mutation. *mrkD* SNP analysis revealed six major *mrkD* variants among the 451 *K. pneumoniae* with 59 SNPS (Figure 5A). In contrast, most of the *treC* were mutated and the variants observed were diverse with 135 SNPs (Figure 5B). However, observed variants of *treC* were strongly associated with the specific STs unlike *mrkD*.

**Figure 5A.**
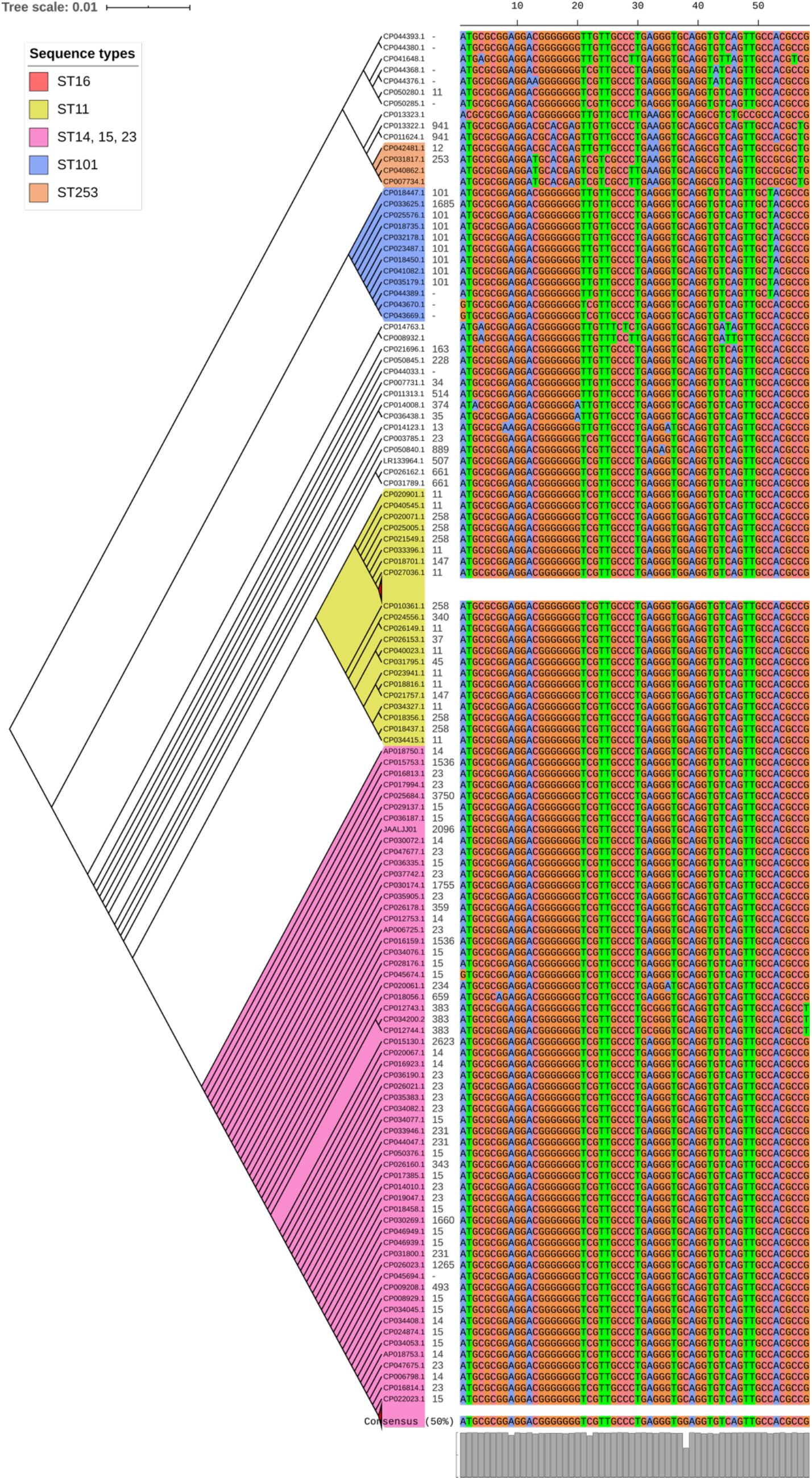
The phylogeny of *mrkD* was based on the maximum-likelihood tree with 59 SNPs. Six major variants were observed as observed in the multi-alignment of SNPs. The STs did not exactly correlate with the variants of *mrkD*, whereas multiples STs share same *mrkD* variants.

**Figure 5B.**
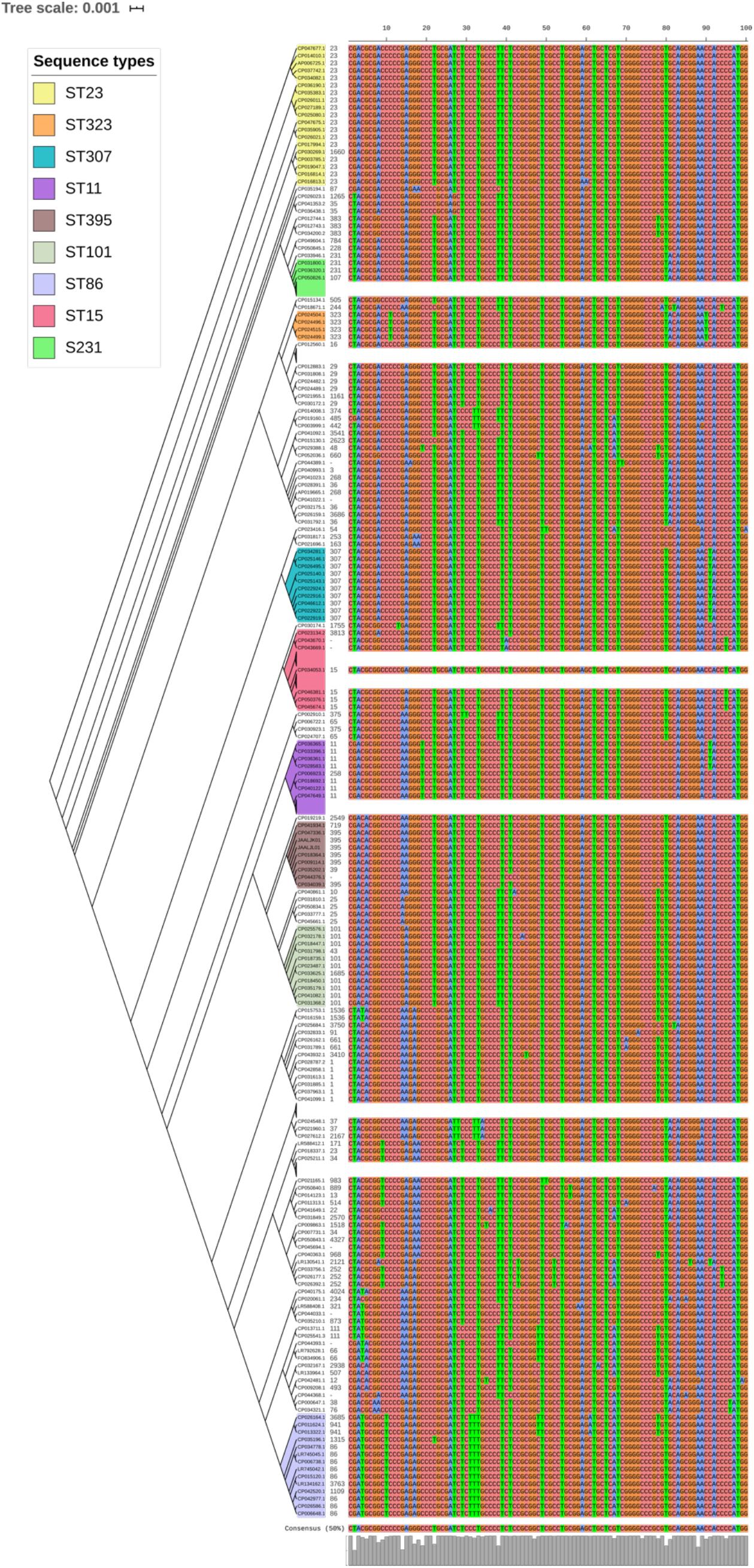
*treC* phylogeny was based on the maximum-likelihood tree with 135 SNPs. Six major variants were observed as observed in the multi-alignment of SNPs. The STs did not exactly correlate with the variants of *mrkD*, whereas multiples STs share same *mrkD* variants.

Unlike *K. pneumoniae*, in *K. quasipneumoniae* each ST was observed to be specific to each country. ST1813 was found in India, ST489 and ST196 were found in USA, whereas ST3870 and ST3876 were found in China. All isolates harboured *allS, iutA, fimH, mrkD, pilQ, ecpA, pgaA, pgaB, pgaC, treC* and *luxS*. ST526 and ST668 lacked the type I fimbriae gene *fimA*, wheres ST1818 from India lacked *bcsA* gene responsible for adhesin. Except one, none of the isolates harboured *wzc* for CPS but harboured *treC* instead. QS2 family gene *luxS* was found in all isolates. Except ST3870, none of the isolates harboured *wcaJ* gene.

For a better understanding of AMR genes across the globe, country-wise distribution of carbapenemases and *mcr* genes is depicted in figure 6. *bla*_KPC_ was the dominant carbapenemase in USA and China, whereas *bla*_NDM_ was more or equal to *bla*_OXA-48_ like genes in UK, Norway, Sweden, Japan and Korea. In contrast, among India and Europe *bla*_OXA-48_ like was common. In Australia, both *bla*_IMP_ and *bla*_KPC_ were observed. Interestingly, *mcr* genes were observed in China, Taiwan, Thailand and Vietnam. Distribution of plasmids among *K. pneumoniae* are depicted in figure 7. A total of 1286 plasmids were harboured in 454 isolates with 1819 replicon types. All six sequenced isolates commonly harboured IncFIB(K)_Kpn3 plasmids in addition to other replicon types.

**Figure 6.**
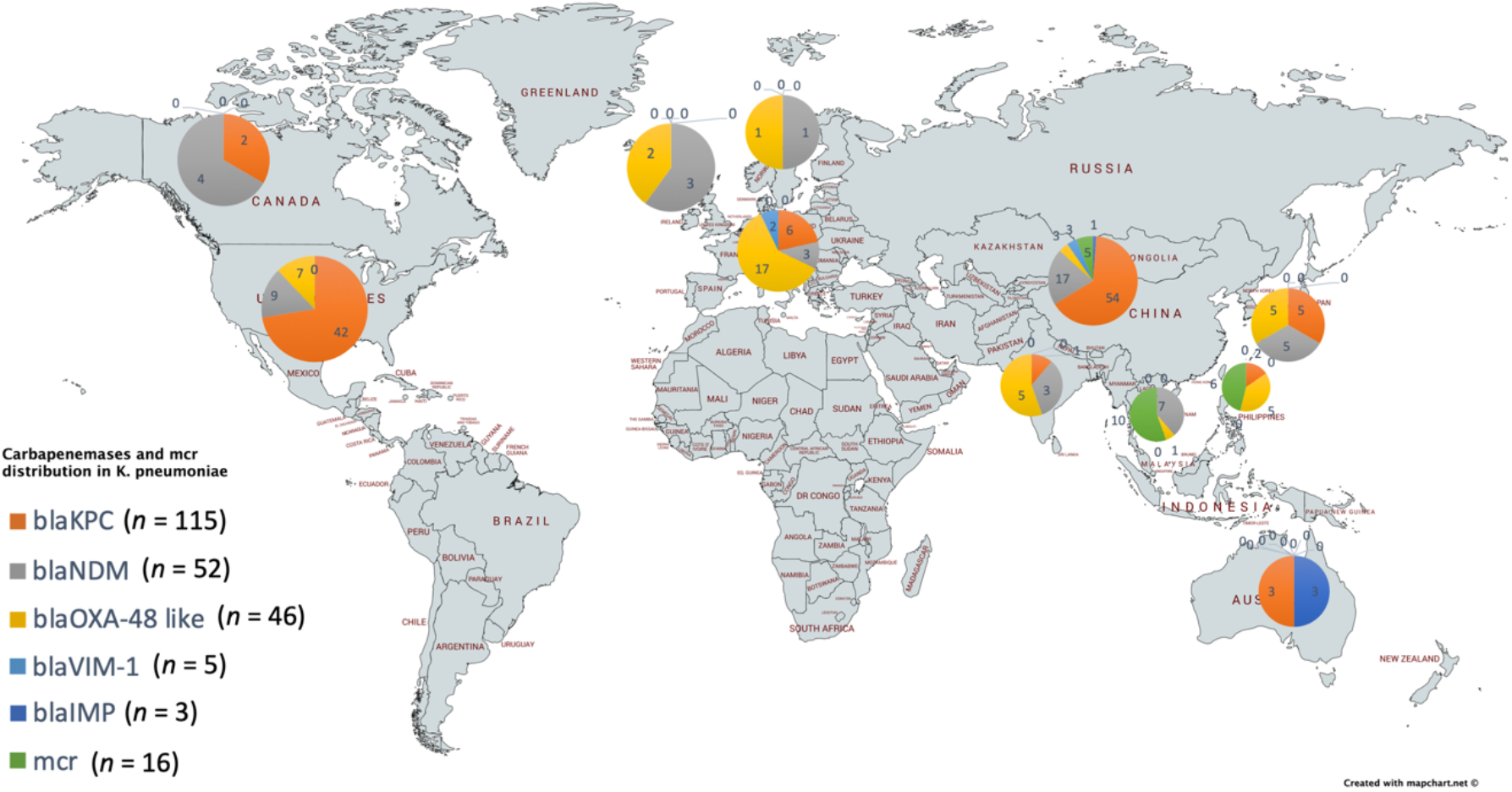
Country-wise distribution of carbapenemases and *mcr* genes in *K. pneumoniae*. Map outline was created using mapchart.net (https://mapchart.net/)

**Figure 7.**
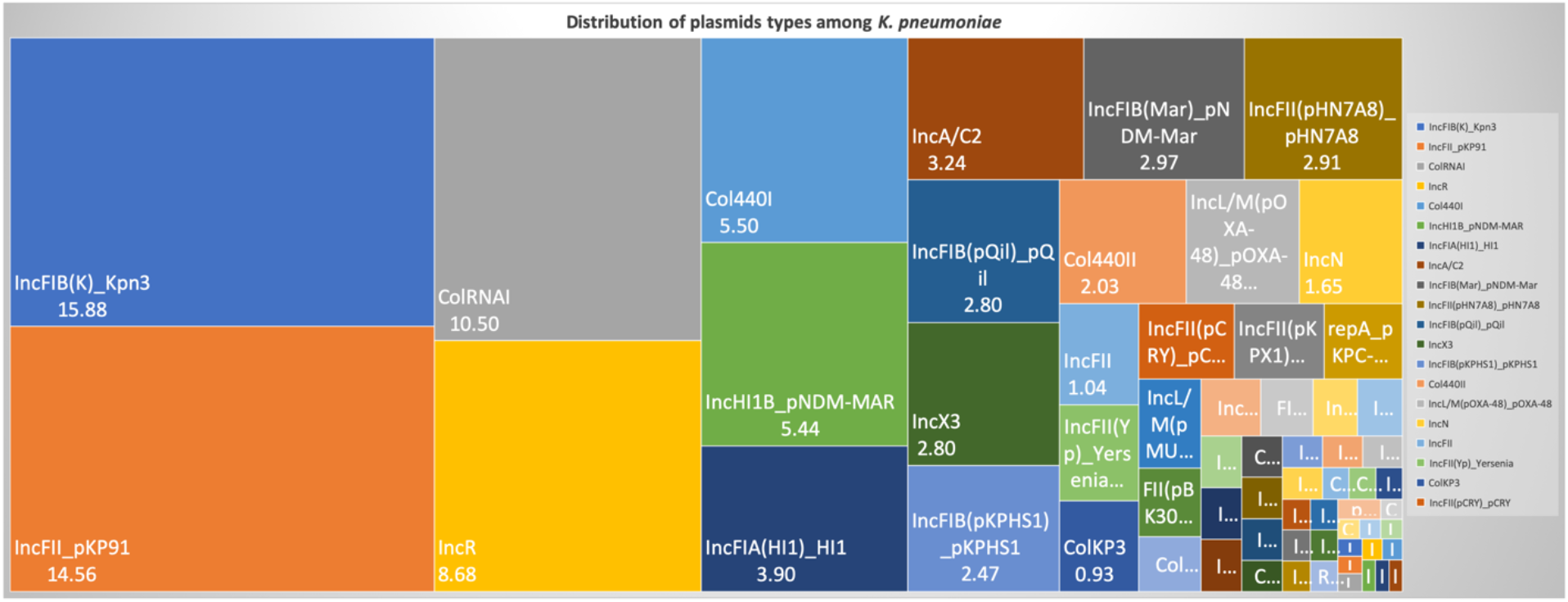
Distribution of plasmid types (percentage) among 454 *K. pneumoniae*. A total of 1286 plasmids were harboured in 454 isolates with 1819 rep types. Plasmids harbouring single replicon type were 791, two replicons - 457, three replicons - 34, and four replicons were 3 plasmids.

Moreover, the association of genetic biofilm factors and AMR genes were analysed using scoary (Table 3). Genes observed significant (p<0.05) with Benjamini-Hochberg’s p-value were only considered. *bla*_IMP_ and *blaVIM* did not associate with any of the screened biofilm genetic factors, whereas *bla*_OXA-48_ like, *bla*_NDM_ and *bla*_KPC_ had significant associations with biofilm genes co-carried with them.

**Table 3:** Association of biofilm genetic virulence factors with antimicrobial resistance genes for carbapenem resistance as analysed using scoary

|  |  | p-values |
| --- | --- | --- |
| *bla<sub>OXA-48</sub>* like | *mrkD*<sup>+</sup> | 0.01** |
|  | *BcsA* | 0.08 |
|  | *WcaJ* | 0.05* |
|  | *pgaA*<sup>+</sup> | 0.05* |
| *bla<sub>NDM</sub>* | *mrkD*<sup>+</sup> | 0.03* |
|  | *fimA* | 0.01** |
|  | *pgaC* | 0.09 |
|  | *wcaJ* | 0.03* |
|  | *iutA* | 0.04* |
| *bla<sub>KPC</sub>* | *mrkD* | 0.03* |
|  | *fimA* | 0.06 |
|  | *fimH* | 0.03* |
|  | *treC* | 0.09 |
|  | *bcsA* | 0.02* |
| *mcr* | *bcsB* | 0.002** |
|  | *iutA* | 0.05* |

<sup>+</sup>hypothetical proteins were significantly co-carried with these biofilm genetic factors
\*p<0.05; \*\*p<0.01

## Discussion

The biofilm-forming capacity of clinical *K. pneumoniae* isolates had a significant association with the outcome in respective patients. Strong biofilm formation significantly reduced the number of days alive for the patient to 3.33 days from the poor/negative biofilm producing isolates with 11.33 days. This clearly demonstrates the effect of biofilm formation as an association for increasing/accelerating the mortality. This is of particular concern, as *K. pneumoniae* can survive for prolonged periods in the hospital environment and grow attached to inanimate surfaces like medical devices (catheters, ventilators)^[2]^.

In such situations, polymicrobial infections might further complicate the therapy. Monomicrobial or polymicrobial isolates were identified in addition to *K. pneumoniae* among 45 out of 72 patients. This includes, *P. aeruginosa*, *Acinetobacter spp.*, *Enterococcus spp.*, *S. aureus*, and *Candida spp*. Almost all of these species were previously recorded in literature for their biofilm-forming abilities. Polymicrobial infections are known to play an inevitable role in managing biofilm structure and causing persistent infection. The polymicrobial infections observed in the study indicate the complexity of antimicrobial therapy and in the selection of appropriate antimicrobials for the patient.

The biofilm structure has been known to fuel/protect the pathogen even in stressful conditions *in vivo*, thereby resulting in clinical failure irrespective of antimicrobial susceptibility. This was clearly reflected in the 35% (*n* = 5) of clinical failure among the carbapenem-susceptible infections (*n* = 14); three of the five clinical failures were strong or moderate biofilm formers. Of the remaining two, one patient had diabetes mellitus as a comorbid condition while the other did not show any co-morbid condition. This clinical failure might be due to a hypervirulent condition that needs further analysis for confirmation. The MBEC assay also confirms the findings, as C3, C5, C6 and C8 strains showed an increase from MICs of ≤ 0.03 μg/ml (planktonic cells) to 2, 32, 128 and 4 μg/ml, respectively for biofilms. These susceptibility results of carbapenem on the fully-grown biofilm structure were consistent with the previous reports for gentamicin, cefotaxime and ciprofloxacin, amikacin and piperacillin in *K. pneumoniae*^[18,19]^. MBEC assays further revealed that the planktonic cells released from biofilm structure exhibited the same MICs as the biofilm cells, which might possibly be a mechanism for establishment of mobile resistant population *in vivo* causing clinical failure.

Treatment failure was observed in a proportion of patients who received carbapenem monotherapy due to development of resistance, particularly in those infected with biofilm forming strains. This shows that biofilm producing strains develops resistance rapidly upon antibiotic exposure and thus lead to addition or change of antimicrobial agent. For CR *K. pneumoniae*, colistin in combination with meropenem was prescribed. Further, for carbapenem-colistin resistant *K. pneumoniae*, aminoglycosides or tigecycline was added in combination for therapy.

Several scientific investigations have hypothesized various physical mechanisms involved in promoting AMR due to biofilm formation. One of the commonly proposed methods has been the ability of biofilm matrix to prevent efficient diffusion of antibiotics, leading to significantly decreased exposure of bacteria in biofilms, in addition to development of resistance due to altered gene expression of antibiotic tolerance genes, and horizontal transfer of AMR genes between cells in a biofilm environment^[20,21]^. In *K. pneumoniae*, it was reported that ampicillin was steadily degraded as it diffused through biofilm matrix. Ciprofloxacin could penetrate the biofilm but could not kill the bacterial cells as they become tolerant at nutrient limited conditions^[22,23]^. In addition, antibiotics piperacillin, piperacillin-tazobactam, cefoperazone, ceftazidime, cefepime, meropenem, ciprofloxacin, netilmicin and amikacin were proven to exhibit reduced activity against adherent bacteria when compared to the planktonic counterparts^[24]^.

The genetic mechanisms known to be involved for a successful biofilm formation in *K. pneumoniae* include factors for adhesion (fimbriae and pili)^[6]^, cohesion (adhesins, polysaccharides), CPS^[7]^, QS^[9]^, and loss of mucoidal nature (colonic acid)^[8]^. Here, genes responsible for each of these factors were analysed for the study isolates in comparison with global clones. Though a few reports suggest recombination between *K. variicola* and *K. quasipneumoniae*^[25]^, and between *K. quasipneumoniae* and *K. pneumoniae*^[26]^, most of the studies report that homologous recombination does not occur between strains in the different clades^[27]^. The phylogenetic analysis in the present study revealed significant differences between *K. pneumoniae* and *K. quasipneumoniae* and hence these were studied separately for further analysis.

Fimbriae and pili structures are known for their ability to support motility as well as attachment to a biotic or an abiotic surface, essential in the initial stages of biofilm formation. The two fimbrial systems involved in *K. pneumoniae* and *K. quasipneumoniae* for these functions are type I and type III fimbrial systems. *fimA* and *fimH* genes are known to be responsible for type I fimbrial system in *K. pneumoniae* whereas the type III fimbriae in *K. pneumoniae* encoded by the *mrk* gene cluster (*mrkABCDF*) contain the structural and assembly components. However, the adhesive property for efficient attachment is conferred by MrkD protein at the tip of the fimbriae, which helps in better attachment to the basolateral surfaces *in vivo* such as urinary tract or bronchial epithelia^[2]^. *mrkD* was proven to play a major role during pathogenesis in a nosocomial infection by facilitating attachment to damaged mucosal surfaces caused due to insertion of indwelling devices like catheters, as well as adherence to the surfaces of these devices which would be coated with host-derived conjugates^[28,29]^. Interestingly, the genes encoding the production of type III fimbriae of *K. pneumoniae* could be either chromosome or plasmid-borne^[30]^. Investigations also reported that the plasmid-borne type III fimbriae gene cluster is associated with a conjugative plasmid that may also be involved in horizontal transfer of AMR genes^[2]^.

Initially, it was considered that type I fimbriae are not as important as type III fimbriae for biofilm formation. However, recent investigations have indicated that either type I or type III fimbriae may play a role in biofilm formation^[31]^. Accordingly, in the present study all six sequenced isolates harboured genes responsible for both type I (*fimA* and *fim*H) and type III (*mkr*D) fimbriae. Also, these genes were present irrespective of sequence types across all global clones.

Several allelic variants of *mrkD* have been identified in different *Klebsiella* isolates. These alleles are known to be associated with different binding specificities to matrix material such as collagen^[32]^. Mutational analysis among the 454 *K. pneumoniae* revealed only six major variants of *mrkD* exhibiting it as a stable genetic factor for biofilm mechanism.

The *E. coli* adhesive structures common pilus (*ecp*) and *pilQ* are known to be important in regulation of pili function in *K. pneumoniae*. In the present study, *ecpA* and *pilQ* were observed in almost all isolates of *K. pneumoniae* and *K. quasipneumoniae*. Similarly, Alcántar-Curiel et al.^[6]^ reported that 96% of the *K. pneumoniae* strains contained *ecpA* with a 94% phenotypic correlation of ECP production during adhesion to cultured epithelial cells, which was justified as a critical factor in forming an adhesive structure. Among *K. pneumoniae* sequenced in this study, one ST2096 (CC14) was negative for *ecpA*, while *pilQ* was absent only in two of the *K. pneumoniae* belonging to ST395. In *K. quasipneumoniae*, all three isolates harboured both *ecpA* and *pilQ* genes.

Although genes responsible for adhering structure were shown to be present, the actual secretory polysaccharides and other adhesins play a critical role in physical attachment that enhances biofilm formation. *pgaABCD* and *bcsA* were highly reported adhesins in *K. pneumoniae*^[33,34]^. The adhesin gene profile correlated with the particular STs. Accordingly, ST11, ST15, ST14 and ST16 carried all four genes; ST258 and ST307 carried *pgaA*, *pgaC* and *bcsA*; ST23 carried *pgaB*, *pgaC* and *bcsA*; while ST101, ST147, ST231 and ST395 carried only *pgaC* and *bcsA*. Chen and co-workers^[33]^ previously reported that the loss of *pgaC* affected the production of poly-β-linked N-acetylglucosamine which in turn had inhibitory effects on *in vitro* biofilm formation. Among the six sequenced isolates, none of them harboured *bcsA*. *K. quasipneumoniae* harboured all three *pga* gens, while in *K. pneumoniae* two isolates harboured *pgaC* and one harboured *pgaB* and *pgaC* genes.

Lipopolysaccharides are known to be involved in the initial adhesion on abiotic surfaces and capsule plays a critical role in construction of mature biofilm architecture^[35]^. *wzc*, *cpsD* and *treC* have been reported to be key regulators in the CPS production^[7]^. Since ST23 were the only isolates found to harbour *wzc* genes in the study population, they were further analysed for their efficiency in hyper-capsulation due to a 565 glycine-to-serine substitution in *wzc* genes. None of the ST23 isolates carried Gly-565Ser substitution, known for hyper-capsulation. ST23 had clearly demonstrated hypervirulence characteristics and harboured genes exclusively for those characteristics such as *wcaG* and *rmpA*/*A2*, although they might not be good biofilm producers. A recent study by Ernst^[36]^ revealed that isolates that did not carry the capsule transcription factors *rmpA/A2* for hypervirulence but carried a mutated *wzc* gene, resulted in hyper-capsulation. This proved that disruption/absence of *wzc* might cause the hyper-capsulation required for biofilm formation irrespective of *rmpA/A2*.

*treC* was found to aid in trehalose utilization through production of trehalose-6-phosphate hydrolase thereby modulating the CPS production. In a study by Wu et al.^[7]^ it was proven that addition of glucose to the culture medium restored the capsule production and biofilm formation in the *treC* mutant. *treC* was identified in almost all study isolates of *K. pneumoniae* and *K. quasipneumoniae*. A separate analysis on *treC* mutations revealed that most of isolates observed to be mutated to *treC* which might impair biofilm formation. Though, there was a correlation between the STs and SNPs within *treC* gene; the variants observed were diverse. Moreover, variants of *mrkD*, *wzc* and *treC* genes in these isolates requires further gene knock-out studies to analyse the effect of these variants among clinical *K. pneumoniae*.

Furthermore, factors well known for hypervirulence, such as *allS*, *iutA*, *rmpA*/*A2*, *magA*, *K2A*, *wabG*, *wcaG* were also reported for their role in biofilm formation^[10,11]^ and *magA*, *K2a*, *rmpA*/*A2*, *wabG* and *wcaG* were shown to regulate CPS synthesis. In fact, *wcaG* was proven to be an independent risk factor for biofilm formation and highly associated with ST23^[10]^. In line with this, these factors were also identified in isolates analysed in this study. *wcaG* and *magA* were exclusive to ST23, whereas *rmpA* and *rmpA2* were seen majorly in ST23, in addition to C4 (ST2096) and other few STs. Almost all *rmpA/A2* observed were plasmid mediated.

QS is a well-established mechanism in the process of biofilm formation, where the population is regulated on sensing the cell-cell contact^[9]^. Previously a Type II QS system was identified in *K. pneumoniae*, which showed the role of *luxS* in autoinducer AI-2 synthesis^[37,38]^. It was also noted that changes in biofilm architecture were observed in the *luxS* mutant *K. pneumoniae* with less surface coverage and reduced macrocolony formation^[9]^. *luxS* was present in almost all isolates in the study irrespective of ST.

A recent study by Pal et al.^[8]^ reported that loss or disrupted *wcaJ* (glycosyltransferase) in *K. pneumoniae* made the isolates less mucoidal and higher in biofilm forming efficiency. Mutant *wcaJ* known to result in the absence of colanic acid rendering a non-mucoid phenotype that was identified to be associated with reduced susceptibility towards both polymyxins and macrophages (less immunogenic). The analysis indicated that except ST3870, none of the *K. quasipneumoniae* harboured *wcaJ* gene. While in *K. pneumoniae*, *wcaJ* was restricted to ST14, ST23, ST147, and part of ST11. Interestingly, ST15, ST16, ST307 and ST258 – reported global high-risk clones were *wcaJ* negative indicating the high potential of biofilm forming capacity in these STs. Sequenced isolates did not harbour *wcaJ* except C1 (ST395).

Some of the previous studies report association of phenotypic biofilm forming efficiency with the antimicrobial resistance genes in *K. pneumoniae*^[39]^. In clinical *K. pneumoniae*, it was reported that among 44.7% of biofilm formers, 45.3% were ESBLs producers^[40]^. However, there was a lacuna of genetic factors being compared for biofilm mechanism with such AMR genes. Results observed in this study clearly exhibit the strong association of genes responsible for biofilm formation and antimicrobial resistance. Though, *bla*_IMP_ and *bla*_VIM_ did not associate with any of the screened biofilm genetic factors, it was evident that *bla*_OXA-48_ like, *bla*_NDM_ and *bla*_KPC_ had significant (p<0.05) associations with biofilm genes. In addition, hypothetical proteins with significant (p<0.05) associations alongside *mrkD* and *pgaA* were identified. Further gene knockout studies on these hypothetical proteins will reveal their role in regulating biofilm formation.

Limitations of the study include, that the study did not include samples directly from biofilms in these patients and retrospectively used isolates obtained from blood or endotracheal aspirate for biofilm screening. Polymicrobial infections should be studied to see the effect of each species in the contribution of biofilm formation. Also, the study included only complete genomes to generate a high-quality phylogeny output so as to avoid bias in the core vs accessory genes being characterised which occurred with the shot-gun genome sequences (data not shown). The small number of complete genomes available in the few STs included in the analysis significantly limited the ability to arrive at definitive conclusions on certain gene correlations.

A single genetic factor is not always adequate, rather it requires a set of genetic factors to facilitate the complete formation of biofilm. Accordingly, this study discussed various genetic factors involved in biofilm formation of *K. pneumoniae* and *K. quasipneumoniae*, which cluster to provide a combined effect called biofilm. These results highlight the importance of biofilm testing, especially for nosocomial infections that are difficult to clear *in vivo*. These infections require additional treatment that might effectively help in improving patient outcome in nosocomial infections due to *K. pneumoniae*. Further, information on the clonal spread of biofilm forming *K. pneumoniae* across the globe will help to understand the dynamics of biofilm infections thereby paving way for effective management of nosocomial infections.

